# The human claustrum tracks slow waves during sleep

**DOI:** 10.1101/2024.01.29.577851

**Authors:** Layton Lamsam, Mingli Liang, Brett Gu, George Sun, Lawrence J. Hirsch, Christopher Pittenger, Alfred P. Kaye, John H. Krystal, Eyiyemisi C. Damisah

**Author notes:** <u>Corresponding Author</u>: Eyiyemisi Damisah, MD, 800 Howard Ave, New Haven, CT 06519.

## Abstract

Slow waves are a distinguishing feature of non-rapid-eye-movement (NREM) sleep, an evolutionarily conserved process critical for brain function. Non-human studies posit that the claustrum, a small subcortical nucleus, coordinates slow waves. We recorded claustrum neurons in humans during sleep. In contrast to neurons from other brain regions, claustrum neurons increased their activity and tracked slow waves during NREM sleep suggesting that the claustrum plays a role in human sleep architecture.

## Main Text

Slow waves are low-frequency (0.3 – 1.5 Hz) electrographic brain oscillations present across the N2 and N3 stages of non-rapid-eye-movement (NREM) sleep and are a defining feature of the deeper N3 stage (known as slow-wave sleep, SWS).^1–4^ NREM sleep is important for brain homeostasis and memory consolidation, and its disruption has been linked to a variety of neurocognitive and sleep disorders.^5^ At the single neuron level, slow waves are characterized by alternating periods of synchronous silence (hyperpolarized DOWN states) and firing (depolarized UP states) in cortex.^6^ Despite great interest in the contribution of slow waves to normal sleep physiology, the source of their regulation is poorly understood in humans.^1,2^

The claustrum is an evolutionarily conserved subcortical sheet of neurons with a high density of reciprocal connections with the cortex.^7–9^ Its ability to modulate many cortical regions simultaneously has motivated inquiry into its diverse functions such as perception, salience detection, and memory.^10–14 11,15,16^ Recent work in reptiles and rodents has shown that the claustrum coordinates slow-wave generation in the cortex during sleep.^15,17–19^ Specifically, the claustrum is thought to regulate slow waves by activating a network of cortical parvalbumin-expressing interneurons, suppressing activity across broad regions of cortex through feedforward inhibition.^11,15,16^ On a longer timescale, claustrum neurons in rodents increase spiking activity during NREM sleep compared to rapid-eye-movement (REM) sleep or wakeful states.^19^ While this function of the claustrum appears to be conserved across reptiles and rodents, it has not yet been observed in humans. We therefore sought to examine the relationship of the human claustrum with slow waves during sleep.

The small axial cross-section of the claustrum and its proximity to the insula and striatum have made human lesion studies difficult to interpret, as lesions frequently entail damage to adjacent structures.^20^ Similarly, electrical stimulation using depth electrodes may not be confined to the claustrum alone, limiting its ability to define claustrum function.^21,22^ To overcome these technical challenges, we robotically placed 40uM microwires into the claustrum of two human subjects with epilepsy to record claustrum neurons. Accurate placement was verified by fusion of pre-operative and post-operative imaging, which projected microwire locations onto subject-specific anatomy **(Fig. 1a-c, Supplementary Fig. 1)**. To identify slow waves, we adapted an established slow-wave detection algorithm for use with field potentials sampled from macroelectrodes in numerous brain regions **(Fig. 1d-f, Supplementary Fig. 2)**. After validating microwire placement and slow-wave detection, we tracked claustrum neuronal activity during periods of slow waves in NREM sleep. Over four nights and 33 hours of sleep, we recorded 49 single units in the claustrum (CLA) along with 73 units in cortical (anterior cingulate cortex, ACC) and subcortical (amygdala, AMY) regions. Simultaneously, we recorded slow-wave activity (SWA) and slow waves (SWs) in 150 intracranial macroelectrode contacts across 77 unique brain regions.

**Figure 1.**
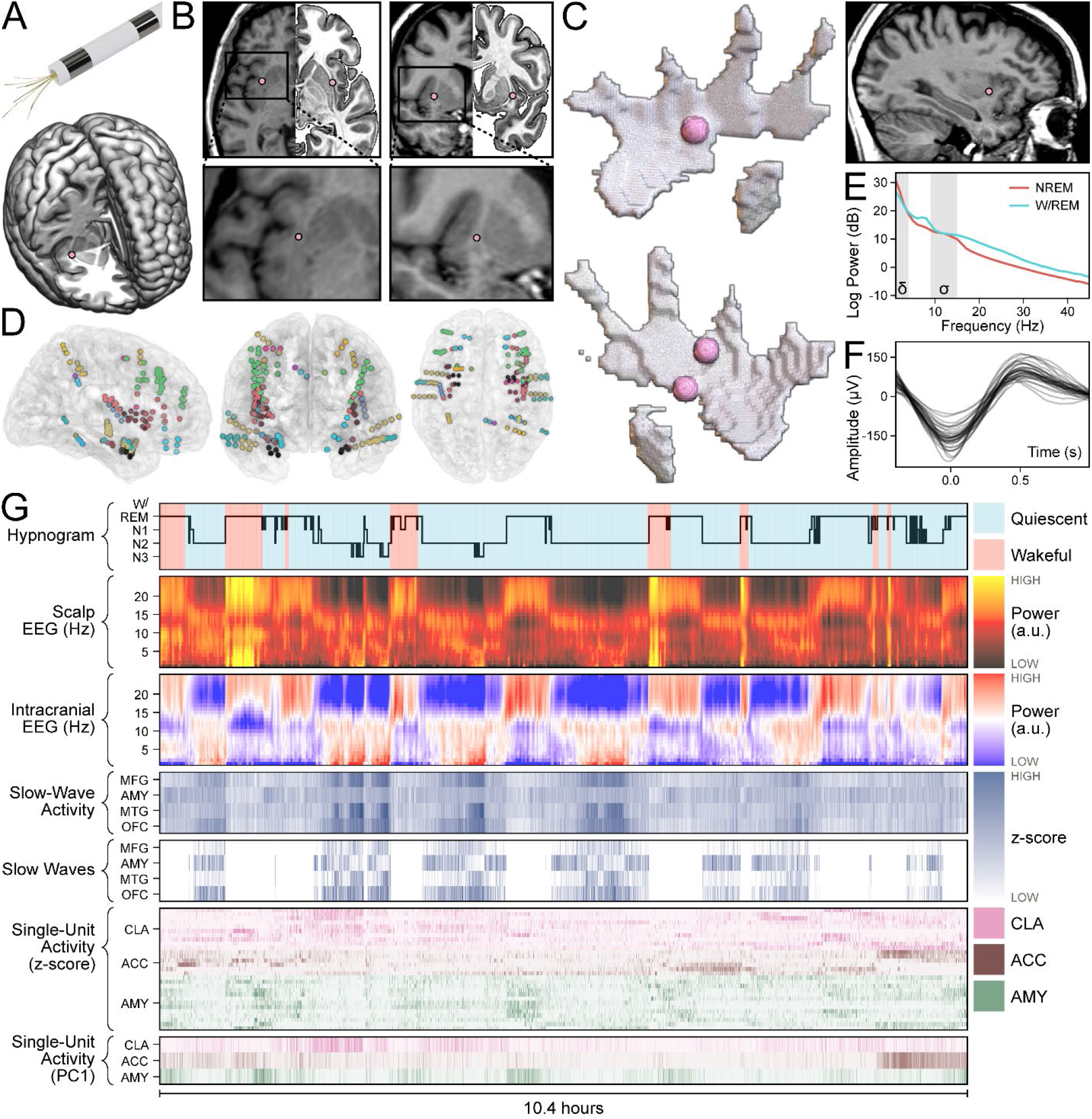
Claustrum single units and cortical slow waves are tracked over hours of sleep. **(a)** Model of a Behnke-Fried depth electrode with protruding microwires for sampling the claustrum (top). Location of right claustrum microwires (pink dot) in Subject A on the MNI152 template (bottom). **(b)** Axial and coronal T1 MR images of Subject A with corresponding mirrored MNI152 templates (top row) marking the location of right claustrum microwires and accompanied by magnified inserts (bottom row). **(c)** Locations of distal microwires when superimposed onto MNI152 models of the right and left claustrum, respectively (left top and bottom). Sagittal T1 MR image of Subject A (top right) marking the location of the right claustrum microwires. **(d)** Electrode locations for all subjects superimposed onto the N27 template in axial, coronal, and sagittal views (left, middle, right). **(e)** Power spectral density of NREM and W/REM sleep across all channels in Subject B, Night 03. Delta and sigma frequency bands (indicating SWA and sleep spindles, respectively) are shaded. **(f)** Average waveforms of detected slow waves in each channel of Subject B, Night 03. **(g)** Sleep recording from Subject B, Night 03. Hypnogram is colored by behavioral state observed on avEEG: red indicates wakefulness and blue indicates behavioral quiescence (first row). Power spectrogram from the C4 scalp electrode (second row). Illustrative power spectrogram from right middle frontal gyrus (third row). Binned z-score of slow-wave activity from four regions: middle frontal gyrus, amygdala, middle temporal gyrus, and orbitofrontal cortex (fourth row). Binned z-score of slow wave presence from the same regions (fifth row). Binned z-score of the firing rate for claustrum (pink), anterior cingulate cortex (brown), and amygdala (green) single units (sixth row). First principal component of the firing rate for single units in the above regions (seventh row).

**Figure 2.**
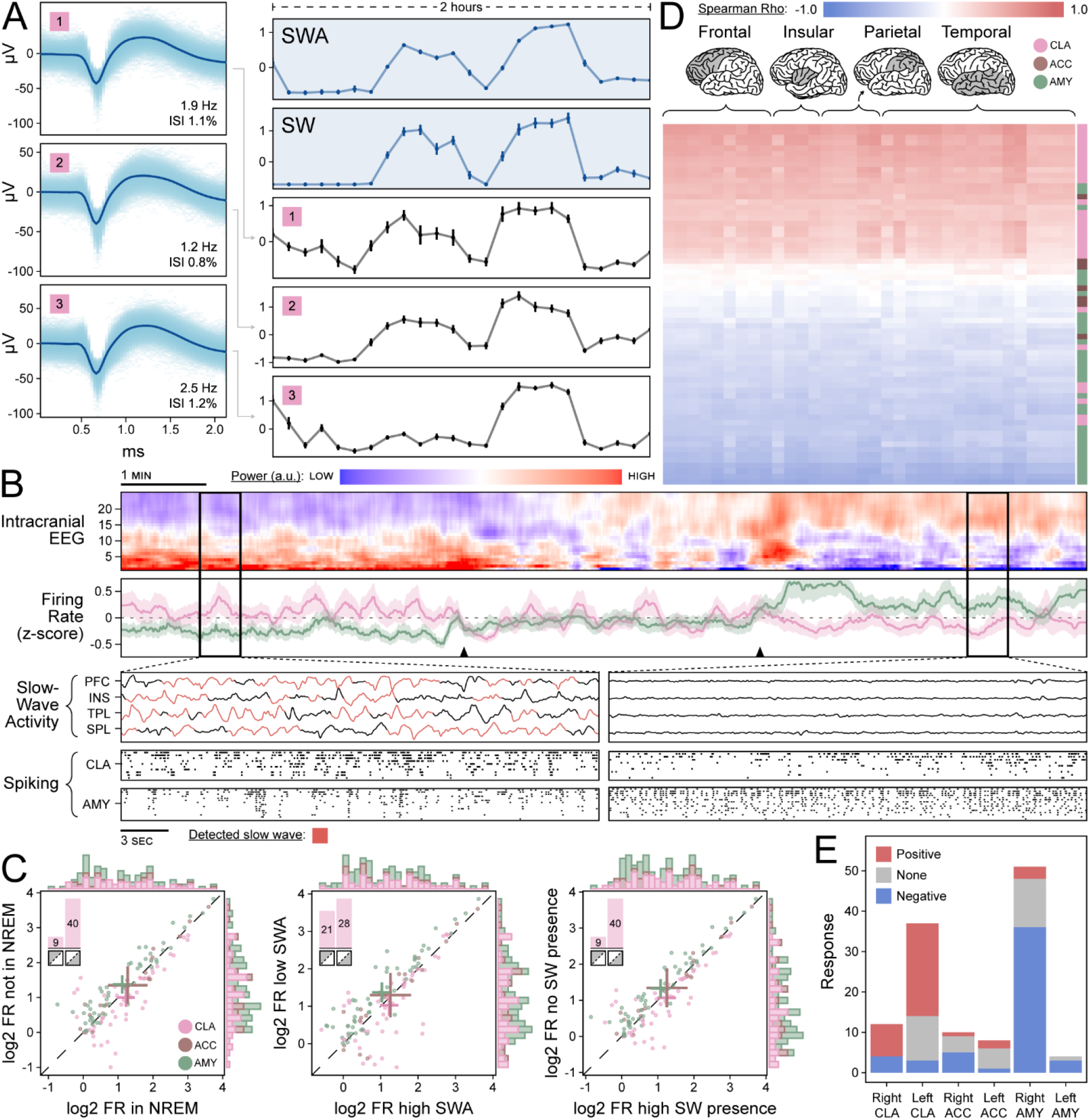
Claustrum single unit activity correlates with slow waves. **(a)** Average waveforms of three claustrum units with inset firing rate and inter-spike interval violations (left panels) with corresponding z-scored firing rate (black panels) aligned to z-scored slow-wave activity (SWA) and slow wave (SW) presence in the right orbitofrontal cortex from Subject A (blue panels) (right panels). Standard error is indicated by vertical bars. **(b)** Transition out of NREM sleep in Subject A. Power spectrogram of left middle frontal electrode (first row). Z-scored population firing rates in the claustrum (pink) and amygdala (green) with black triangles indicating transition period (second row). Magnified windows before and after sleep transition (bottom left and bottom right panels) showing SWA (red highlighting indicates a SW) in the orbitofrontal cortex, middle temporal gyrus, amygdala, and middle frontal gyrus (first rows of bottom panels) and raster plots of spiking activity for the claustrum (second rows of bottom panels) and amygdala (third rows of bottom panels). **(c)** Scatterplots of log2 firing rate of claustrum (pink), anterior cingulate (brown), and amygdala (green) units stratified by three measures of SWs: sleep stage (not NREM sleep vs. NREM sleep, left), SWA (25^th^ vs. 75^th^ percentiles, middle), and slow-wave presence (no SW presence vs. greater than mean SW presence, right). Crosses indicate population averages with 95% confidence intervals. Units in the lower triangle favor SWs. Inset bar plots indicate the proportion of claustrum units in the upper vs. lower triangles. **(d)** Heatmap of Spearman’s ρ correlations between units (rows) and SWA across channels (columns) for Subject A. Colors indicate claustrum (pink), anterior cingulate (brown), and amygdala (green) single units. Highlighted lobes indicate channel locations. **(e)** Bar plot showing distribution of single units across all subjects with positive (red), negative (blue), or no correlation (gray) with ipsilateral SWA and SWs.

Studies have shown that neurons typically decrease spiking activity during NREM sleep compared to REM sleep or wakefulness (except for neurons with very low firing rates < 1 Hz).^23–25^ However, the opposite pattern has been observed in the claustrum.^15,19^ We therefore hypothesized that single units in the CLA would increase spiking activity during periods of SWs in NREM sleep and that neurons in other brain regions (the ACC and AMY) would concomitantly decrease spiking activity.

We found that a majority (40/49) of CLA neurons increased spiking activity during periods of SWs in NREM sleep over a timescale of hours; in contrast, ACC and AMY neurons decreased spiking activity **(Fig. 1g and 2a, Supplementary Fig. 3 and 4;** data corresponding to illustrative examples can be found in the supplementary information). On a timescale of minutes, this pattern was most prominent during transitions out of NREM sleep in which CLA spiking activity decreased and AMY population activity increased **(Fig. 2b, Supplementary Fig. 5**). These findings were consistent with observations of claustrum activity during NREM sleep in rodents.

**Figure 3.**
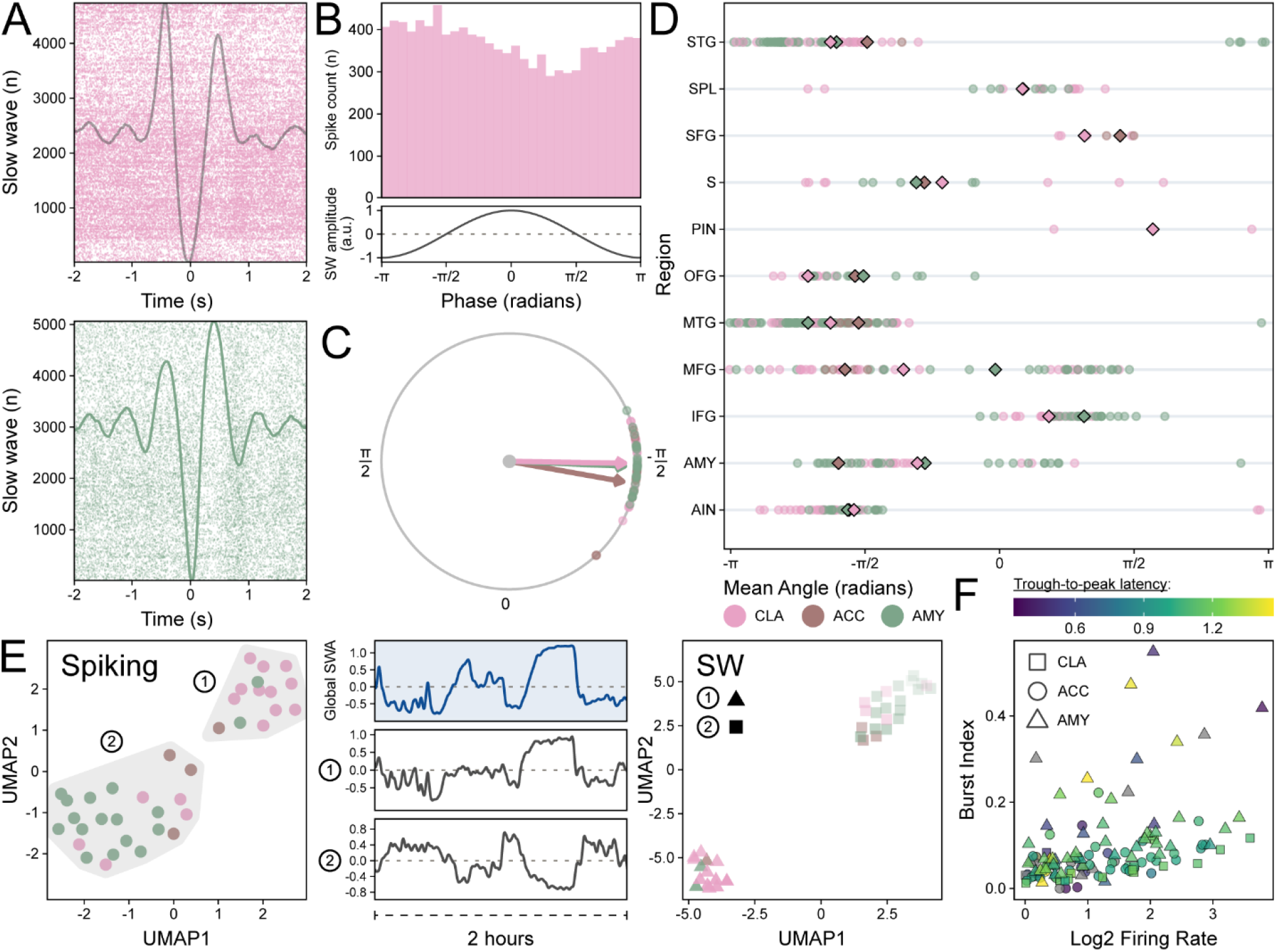
Claustrum phase-locking and population activity correlate with slow waves. (**a**) Raster plot demonstrating phase-locking of an illustrative claustrum unit with slow waves (SW) in an orbitofrontal channel (top) compared to phase-locking in an amygdala unit with its adjacent amygdala channel (bottom) in Subject B, Night 03. The average slow-wave waveforms are superimposed. **(b)** Phase histogram for the same claustrum-orbitofrontal pair (top); the amplitude of an idealized slow wave for each phase is shown (bottom). **(c)** Polar plot for the same claustrum-orbitofrontal pair indicating the preferred phase angle for every unit-orbitofrontal pair with a phase distribution significantly different from uniform. Color indicates unit region, and arrows indicate the average preferred phase angle among all pairs of the same unit region. (**d)** Region-wise preferred phase angles for all unit-channel pairs with a phase distribution significantly different from uniform in Subject A. Diamonds indicate the average of preferred phase angles for each unit region. See **Supplementary Table 1** for abbreviations. **(e)** Scatterplot of UMAP dimensions for Subject A’s single unit spiking activity demonstrating self-segregation of claustrum (pink), anterior cingulate (brown), and amygdala (green) units into two groups (labeled 1 and 2) indicated by gray shading (left). Global slow-wave activity (middle, top panel) with aligned z-scored population firing rates for both groups (middle, middle and bottom panels). Scatterplot of two UMAP dimensions for the same units after dimensionality reduction of correlation with slow wave presence across channels (right). Group 1 units are indicated by triangles, and Group 2 units are indicated by squares. **(f)** Scatterplot of the burst index and log-base-2 firing rate of claustrum (square), anterior cingulate (circle), and amygdala (triangle). Color indicates the trough-to-peak latency in milliseconds.

We then divided sleep recordings into epochs classified by three measures of SWs (sleep stage, SWA, and SW presence). We found that CLA unit activity was significantly higher during periods of SWs – concordant across all three measures – in comparison to the pooled ACC and AMY units (χ^2^: p = 3.0 x 10^-8^, 3.6 x 10^-5^, and 2.9 x 10^-10^, respectively) (**Fig. 2c**). Based on these observations, we quantified the heterogeneity of responses among neurons: Single units were classified into positive-, negative-, and non-responding units based on Spearman’s ρ correlations with SWA and SW in ipsilateral brain regions **(Fig 2d-e, Supplementary Fig. 6)**. This analysis showed that 63% of CLA single units were positive responders compared to 8% of units in other regions (17% ACC, 5% AMY); conversely, only 14% of CLA single units were negative responders compared to 62% of units in other regions (33% ACC, 71% AMY). Thus, the majority of CLA units displayed a preference for spiking with SWs compared to neurons in other regions.

Next, we explored the relationship between the CLA and individual SWs with spike-phase coupling. Significant SW phase locking of CLA units was present in nearly all sampled regions. However, these findings were not unique, as similar SW phase locking relationships were seen with ACC and AMY units **(Fig. 3a-d, Supplementary Fig. 7)**. After characterizing CLA neurons on a timescale of seconds, we collapsed time using Uniform Manifold Approximation and Projection (UMAP) to compare the population activities of the CLA, ACC, and AMY in an unsupervised manner.^26^ Single units segregated into two clusters in UMAP space: One group was enriched in CLA units and positively correlated with SWA, while a second group was enriched in ACC and AMY units and negatively correlated with SWA **(Fig. 3e, Supplementary Fig. 8)**. This grouping was preserved when collapsing the Spearman’s ρ correlations of single units with SWA and SWs, respectively **(Fig. 3e, Supplementary Fig. 8)**. Finally, we inferred the cell types of our 122 single neurons using established metrics **(Fig. 3f)**.^27^ Most CLA units were classified as pyramidal cells (84%) with a similar proportion in control region units (AMY 89%, ACC 83%); the remaining cells were classified as interneurons or did not meet criteria for either cell type.

In this study, we recorded single neurons in the human claustrum for the first time. In contrast to neurons in other brain regions, CLA neuronal activity tracked SWs in NREM sleep at multiple timescales and over multiple nights of sleep: Spiking activity increased with SWs at NREM sleep onset, maintained heightened levels throughout NREM sleep periods, and decreased with the transition to REM sleep or wakefulness. These correlations were robust for SWs in many brain regions including the frontal lobe, which is known to have strong connections with the human claustrum.^9,15,28^ While CLA phase locking with cortical SWs was not unique, its prevalence in sampled cortical regions is consistent with recent animal findings that CLA neuronal stimulation aligns SWs across the cortex.^19^ Our sample of CLA neurons were mostly classified as pyramidal cells; this result may be consistent with the theory of CLA feedforward inhibition, as CLA pyramidal cells are connected to SW-generating cortex in animals.^11,15,16^ In summary, we found that human claustrum spiking activity increases with SWs during sleep, which supports previous work in animals and may reflect a conserved role of the claustrum in the coordination of human SWs.

We studied claustrum activity in subjects with epilepsy undergoing inpatient seizure monitoring, which may reduce the generalizability of our results to the broader population. In addition, while the CLA may be divided into sub-regions with anatomically distinct efferent projections, we could not directly identify the efferent targets of our sampled neurons.^29^ We therefore had to consider all possible connections, reducing the specificity of our claims.

Finally, in accord with animal literature, we found that a population of human claustrum neurons increased their activity with SWs during NREM sleep. This relationship was consistent across multiple timescales and several measures of SWs. Taken together, our observations build on existing causal studies in animals to converge on the theory that the human claustrum plays a key role in coordinating SWs during NREM sleep. These results advance our understanding of the normal physiology of human NREM sleep and suggest that claustrum disruption may lead to sleep-related neuropathology.

## Acknowledgements

We thank the members of the Yale Comprehensive Epilepsy Center for their excellent patient care, the patients who participated in this study, and Dr. Pue Farooque for her assistance in the internal validation of automatic sleep stage classification.

## Online Methods

### Participants

Two right-hand dominant female subjects (Subject A was 47 years old; Subject B 27 years old) with medication-refractory epilepsy undergoing intracranial EEG (icEEG) electrode implantation for clinical seizure localization were enrolled in the study after providing informed consent. Plans for the icEEG studies were made exclusively for clinical purposes; both subjects had macroelectrodes placed into the middle insula, and microwires extending from those electrodes sampled the claustrum (bilaterally in Subject A, left in Subject B). Both subjects also remained on their home anti-seizure medications for the duration of the study, and sleep sessions were recorded in the epilepsy monitoring unit. One sleep session was recorded for Subject A (2 hours) and three sleep sessions were recorded for Subject B (9.7, 10.6, and 10.4 hours). The study was approved by the Institutional Review Board at Yale University.

### Electrophysiology System

Subdermal scalp electrodes were placed according to the 10-20 system. Spencer Probe and Behnke-Fried depth electrodes (Ad-Tech Medical Instrument Corp., Oak Creek, WI, USA) were placed using a ROSA surgical robot (Zimmer Biomet, Warsaw, IN, USA) to bilaterally sample field potentials from numerous brain regions.^1^ Scalp and macroelectrode data were recorded with a NeuroPort Neural Signal Processor (Blackrock Neurotech, Salt Lake City, UT, USA) at a sampling rate of 2048 Hz with a 0.3 Hz – 500 Hz hardware bandpass filter. Extracellular microwire recordings were sampled at 30 kHz with a 250 Hz – 7.5 kHz bandpass filter in Subject A and a 0.3 Hz – 7.5 kHz bandpass filter in Subject B. The online reference electrodes were screwed into the outer cortex of the frontal bone, and left orbitofrontal contacts were selected as the ground electrodes in both subjects.

### Electrode Localization

Pre-operative axial non-contrast T1-weighted magnetic resonance (MR) sequences with 1mm slices were co-registered with post-operative axial non-contrast CT scans with 0.625mm slices using Statistical Parametric Mapping (SPM) 12.^2^ Macro- and microelectrodes were automatically localized on the fused image and manually adjusted using LeGUI software.^3^ Gray and white matter was automatically segmented, and macroelectrodes in white matter were later excluded from analysis after re-referencing. The fused image and electrode positions were warped into Montreal Neurological Institute (MNI) 152 space for assignment of electrodes to the nearest region of interest (ROI) in the Yale Brain Atlas (YBA).^4^ Electrode coordinates were then transformed into MNI305 and then tkrRAS space for visualization in MRIcroGL and the threeBrain package.^5,6^ A total of 76 contacts in Subject A and 74 contacts in Subject B were selected for analysis **(Supplementary Table 2)**.

### Pre-processing

The Blackrock Neural Processing Matlab Kit (NPMK) was used to concatenate data and repair segments of packet loss (https://github.com/BlackrockNeurotech/NPMK). Data were then cropped to sleep intervals, notch filtered for 60 Hz electrical line noise (and its harmonics), lowpass filtered at 128 Hz, and decimated to 256 Hz using the MNE software library.^7^ Subdermal scalp and intracranial macroelectrodes were separately re-referenced to the common average reference (CAR) for their electrode type.

### Sleep Staging

Sleep recordings were divided into 30-second epochs, and a previously validated algorithm with integral artifact rejection was used to automatically classify sleep stages based on scalp EEG (C4 in both subjects).^8^ Automatic sleep staging was internally validated using manual sleep stage classification by a board-certified neurologist. Sleep staging yielded 65 minutes of N2/N3 sleep in Subject A and 240, 319, and 321 minutes of N2/N3 sleep in Subject B’s three recordings **(Supplementary Table 3)**.

### Artifact Rejection

For automatic intracranial macroelectrode channel rejection, the raw time series was converted into a low frequency time series (0 – 50 Hz), high frequency time series (50 – 120 Hz), and first derivative time series. The Local Outlier Factor (LOF) algorithm (10 neighbors) was then computed for each time series, and any channel with an LOF score less than -2 for any time series was rejected. The power spectral density across the entire sleep recording was computed for every channel using Welch’s method, and outliers were manually identified and cross-referenced to confirm that all channels contaminated by artifact had been removed. Next, interictal epileptiform discharges (IEDs) were detected using a previously developed algorithm, and time intervals within one second of an IED were excluded from subsequent slow-wave detection in a channel-wise manner (https://github.com/Kleen-Lab/Linelength-spike-detector-PYTHON). Finally, sleep intervals classified as N2 or N3 sleep were divided into 3-second epochs for further artifact rejection. Numerous features were extracted for each epoch (mean, variance, standard deviation, peak-to-peak amplitude, skewness, kurtosis, root-mean squared value, quantile, zero crossings, Hurst exponent, band-wise average power for eight frequency bands covering 0.5 – 128 Hz, and slope of the power spectral density).^9^ The LOF score was then computed on the extracted features for each epoch in a channel-wise manner, and epochs with an LOF score of less than -2 in more than three channels were excluded from slow-wave detection (with an additional one-second interval before and after the epoch time interval).

### Audiovisual Review

Audiovisual recordings of candidate sleep sessions were manually reviewed at five-minute intervals on a Natus Neuroworks clinical EEG system (Natus Medical Inc., Middleton, WI, USA). Behavioral state was classified as either wakeful or quiescent, and these results were manually compared to the hypnogram generated by the sleep staging algorithm for each candidate sleep session.

### Time-Frequency Processing

Time-frequency power spectrograms were generated with the Morlet wavelet transform (cycles = 6) for 1 – 25 Hz and log-ratio normalized by frequency. Spectrograms for whole-sleep recordings were averaged into 30-second time bins and smoothed with a Gaussian window. Slow-wave activity (SWA) was extracted from the envelope of the Hilbert Transform on the band-passed time series (0.3 – 4 Hz). Sleep transitions were identified via manual review of whole-recording intracranial spectrograms.

### Slow-Wave Detection

A previously validated algorithm for slow-wave detection on scalp EEG was adapted for use on intracranial EEG.^8^ Only artifact-free intracranial macroelectrode contacts were selected for use. By default, data is 0.3 – 1.5Hz bandpass filtered, both negative and positive peaks are labeled and sequentially paired, feature thresholding is applied, and outliers are removed using the Isolation Forest algorithm. For adaptation to icEEG, raw time series were transformed into channel-wise robust z-scores (rZS) before slow-wave detection. Feature thresholds were set for the duration of negative deflection (0.3 – 1.5s), duration of positive deflection (0.1 – 1.5s), negative peak amplitude (> 1 rZS), positive peak amplitude (> 1 rZS), and peak-to-peak amplitude (> 4 rZS). Channels with < 3 detected slow waves per minute across time intervals classified as N2 or N3 sleep were excluded from analysis. After exclusion, we detected a median of 8.0 SWs per contact per minute of N2/3 sleep in Subject A and 6.3, 7.0, and 6.8 in the three Subject B recordings **(Supplementary Figure 9)**.

### Spike Sorting

The Combinato algorithm was used to detect spikes, reject artifacts, and cluster spikes into units using a validated superparamagnetic clustering algorithm (maximum cluster distance for grouping = 2.5, minimum spikes for cluster selection = 25).^10^ Candidate units were manually reviewed for artifacts and split or merged as appropriate. Candidate units with an average firing rate of < 1 Hz or an inter-spike interval < 3ms in more than 5% of spikes were excluded from analysis **(Supplementary Figure 10)**. Spike sorting yielded 49 claustrum single units (37 left, 12 right), 18 anterior cingulate single units (8 left, 10 right), and 55 amygdala single units (4 left, 51 right) across all recordings **(Supplementary Table 4)**.

### SW Measures

Data were divided into 10-second epochs, and for each epoch, three SW measures were calculated: sleep stage (assigned based on 30-second epochs from sleep staging), z-score of the average log-normalized delta power (SWA), and slow-wave presence (percent of the epoch that a slow wave was present). Adjacent slow waves within one second were merged when calculating slow-wave presence. For each SW measure scatterplot, the χ^2^ test was used to compare the proportion of CLA units in the upper and lower triangles of the scatterplot to those of other regions (ACC and AMY).

### Unit Responsiveness

For each unit, the average firing rate was separately compared epoch-wise to each of the three SW measures using a Spearman’s ρ correlation; this comparison was channel-wise in the case of SWA and SW presence. Spearman’s ρ correlation p-values were corrected for the false discovery rate (FDR) using the Benjamini-Hochberg method. Units were defined as positive responders if they had a positive Spearman’s ρ with FDR-corrected p-values < 0.01 across at least half of the ipsilateral intracranial channels for both SWA and SW presence. Negative responders were defined conversely, and units failing to meet either criterion were defined as non-responders.

### Spike-Phase Coupling

The Hilbert transform was used to extract the phase of the 0.3 – 1.5 Hz frequency band for every macroelectrode channel. Spike times of single units during N2 or N3 sleep were then intersected with the nearest sampled phase, and these phases were pooled for every unit-channel pair. The set of phases for each unit-channel pair was tested for a significant difference from the uniform distribution using Rayleigh’s test with FDR correction.

### Dimensionality Reduction

Spiking activity of single units was binned into 10-second epochs across sleep recordings and z-scored. This data was collapsed from an *n* x *t* matrix (where *t* represents time bins) into an *n* x 2 matrix using Uniform Manifold Approximation and Projection (UMAP).^11^ Correlation matrices of the Spearman’s ρ values between single unit activity and either the SWA or SW presence on macroelectrode channels (after binning into 10-second epochs) were also reduced from *n* x *c* matrices into *n* x 2 matrices (where *c* represents the macroelectrode channels) with UMAP. Unit clusters were then manually isolated for further analysis.

### Cell Type Classification

Several established metrics (firing rate, burst index, and trough-to-peak latency) for inferring cell type were calculated for each single unit.^12^ Burst index was defined as the number of spikes occurring within 10ms of each other divided by the total number of spikes, and trough-to-peak latency was not calculated for positive-spiking units. Thresholds favoring classification as a pyramidal cell were firing rate < 10 Hz, trough-to-peak latency > 0.5 ms, and burst index > 0.2. Single units were classified as pyramidal if the majority of criteria favored classification as pyramidal; they were classified as interneurons if the converse was true. In positive-spiking units where trough-to-peak latency could not be calculated, single units with ties in the number of criteria were classified as an unknown cell type.

