## Supplementary Material for "The human claustrum tracks slow waves during sleep"

### Supplementary Tables

| | |
|---|---|
| STG | Superior temporal gyrus |
| SPL | Superior parietal lobule |
| SFG | Superior frontal gyrus |
| S | Somatosensory |
| PIN | Posterior insula |
| OFG | Orbitofrontal gyrus |
| MTG | Middle temporal gyrus |
| MFG | Middle frontal gyrus |
| IFG | Inferior frontal gyrus |
| AMY | Amygdala |
| AIN | Anterior insula |

**Supplementary Table 1.** Abbreviations for Figure 3d.

| | Subject A | | Subject B | |
|---|---|---|---|---|
| | Left | Right | Left | Right |
| Amygdala | 1 | 2 | 3 | 2 |
| Anterior insula | 5 | 6 | 0 | 2 |
| Hippocampus | 1 | 1 | 1 | 2 |
| Inferior frontal gyrus | 1 | 4 | 0 | 5 |
| Middle cingulate gyrus | 1 | 0 | 0 | 0 |
| Middle frontal gyrus | 5 | 8 | 6 | 12 |
| Middle temporal gyrus | 2 | 6 | 2 | 4 |
| Orbitofrontal gyrus | 3 | 2 | 2 | 2 |
| Posterior insula | 4 | 0 | 0 | 0 |
| Somatosensory | 1 | 4 | 1 | 2 |
| Superior frontal gyrus | 3 | 0 | 0 | 0 |
| Superior parietal lobule | 3 | 2 | 0 | 4 |
| Superior temporal gyrus | 7 | 3 | 6 | 6 |
| Premotor | 0 | 1 | 0 | 3 |
| Temporal pole | 0 | 0 | 1 | 0 |
| Inferior temporal gyrus | 0 | 0 | 0 | 1 |
| Motor | 0 | 0 | 0 | 1 |
| Fusiform gyrus | 0 | 0 | 0 | 3 |
| Posterior cingulate gyrus | 0 | 0 | 0 | 2 |
| Precuneus | 0 | 0 | 0 | 1 |

**Supplementary Table 2.** Gray-matter macroelectrode contacts classified by customized regions of interest derived from the Yale Brain Atlas (prior to channel rejection).

| | Subject A Night 01 | Subject B Night 01 | Subject B Night 02 | Subject B Night 03 |
|---|---|---|---|---|
| W | 40 | 181 | 154.5 | 165.5 |
| R | 8.5 | 122.5 | 128 | 104 |
| N1 | 7 | 37.5 | 31.5 | 33.5 |
| N2 | 62.5 | 234 | 303.5 | 305 |
| N3 | 2 | 5.5 | 15 | 16 |

**Supplementary Table 3.** Sleep staging stratified by sleep recording (prior to artifact detection).

| | | Claustrum | | | | Anterior Cingulate | | | | Amygdala | | | |
|---|---|---|---|---|---|---|---|---|---|---|---|---|---|
| | | Left | | Right | | Left | | Right | | Left | | Right | |
| | | - | + | - | + | - | + | - | + | - | + | - | + |
| Subject A | Night 01 | 3 | 1 | 8 | 4 | 1 | 2 | 1 | 0 | 0 | 0 | 10 | 5 |
| Subject B | Night 01 | 12 | 0 | | | 1 | 0 | 3 | 0 | 2 | 0 | 11 | 0 |
| Subject B | Night 02 | 11 | 0 | | | 1 | 0 | 3 | 0 | 1 | 0 | 8 | 0 |
| Subject B | Night 03 | 10 | 0 | | | 3 | 0 | 3 | 0 | 1 | 0 | 10 | 0 |

**Supplementary Table 4.** Number of single units per electrode and sleep recording.

### Supplementary Figures

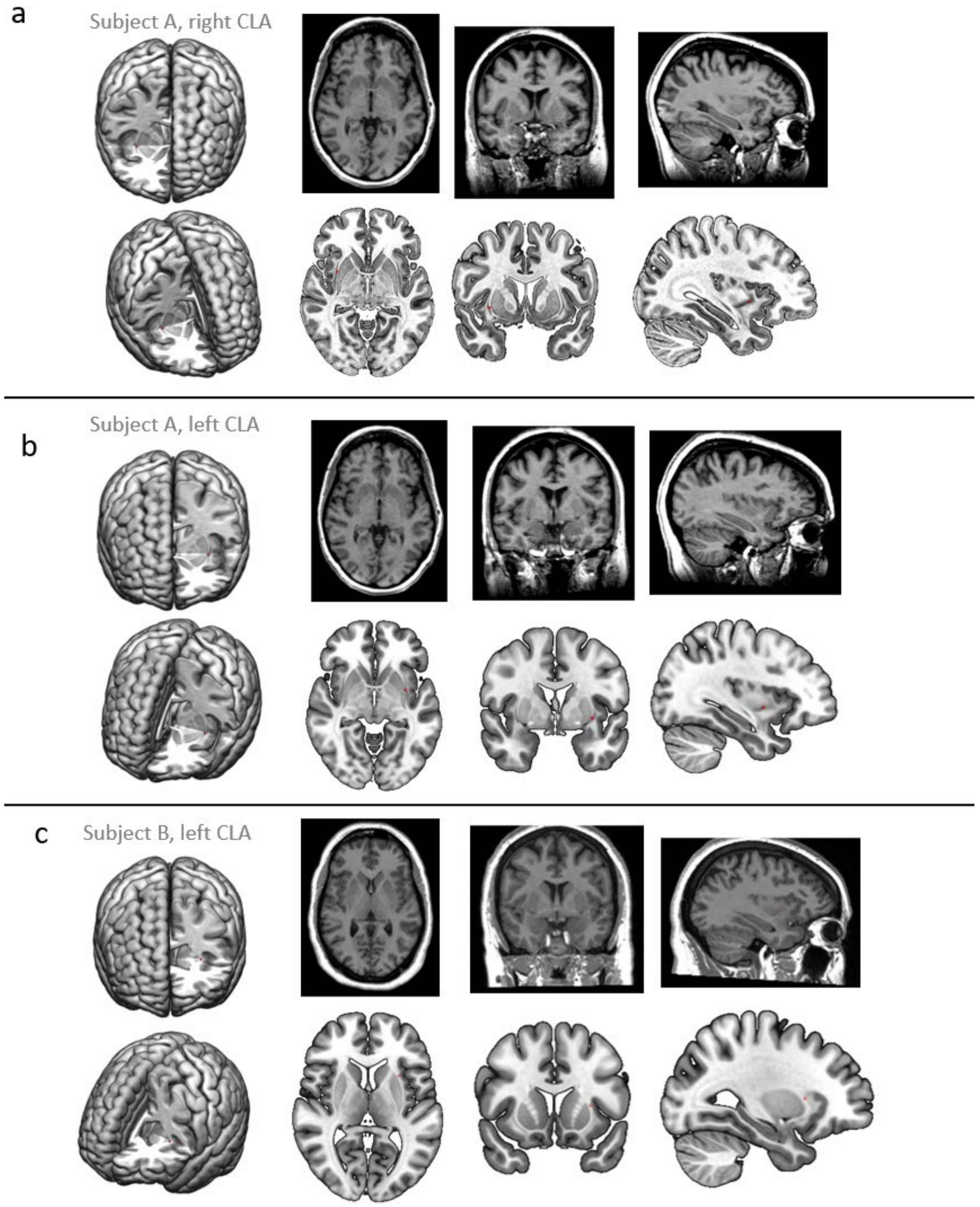


**Supplementary Figure 1. Location of claustrum microwires**. (**a**) Subject A right claustrum (MNI152: x = 32.44, y = 5.98, z = -3.50), (**b**) Subject A left claustrum (-33.15, 0.08, -4.20), and (**c**) Subject B left claustrum (-32.73, 3.51, 1.63). Three-dimensional reconstructions on MNI152 brain models with cutouts to illustrate CLA microwire placement (red dot) in anterior (top left) and oblique (bottom left) views. Placement is also demonstrated by red dots on T1 MRI axial (top left middle), coronal (top right middle), and sagittal (top right) slices; corresponding slices on the MNI152 template shown with a red dot on axial (bottom left middle), coronal (bottom right middle), and sagittal (bottom right) slices.

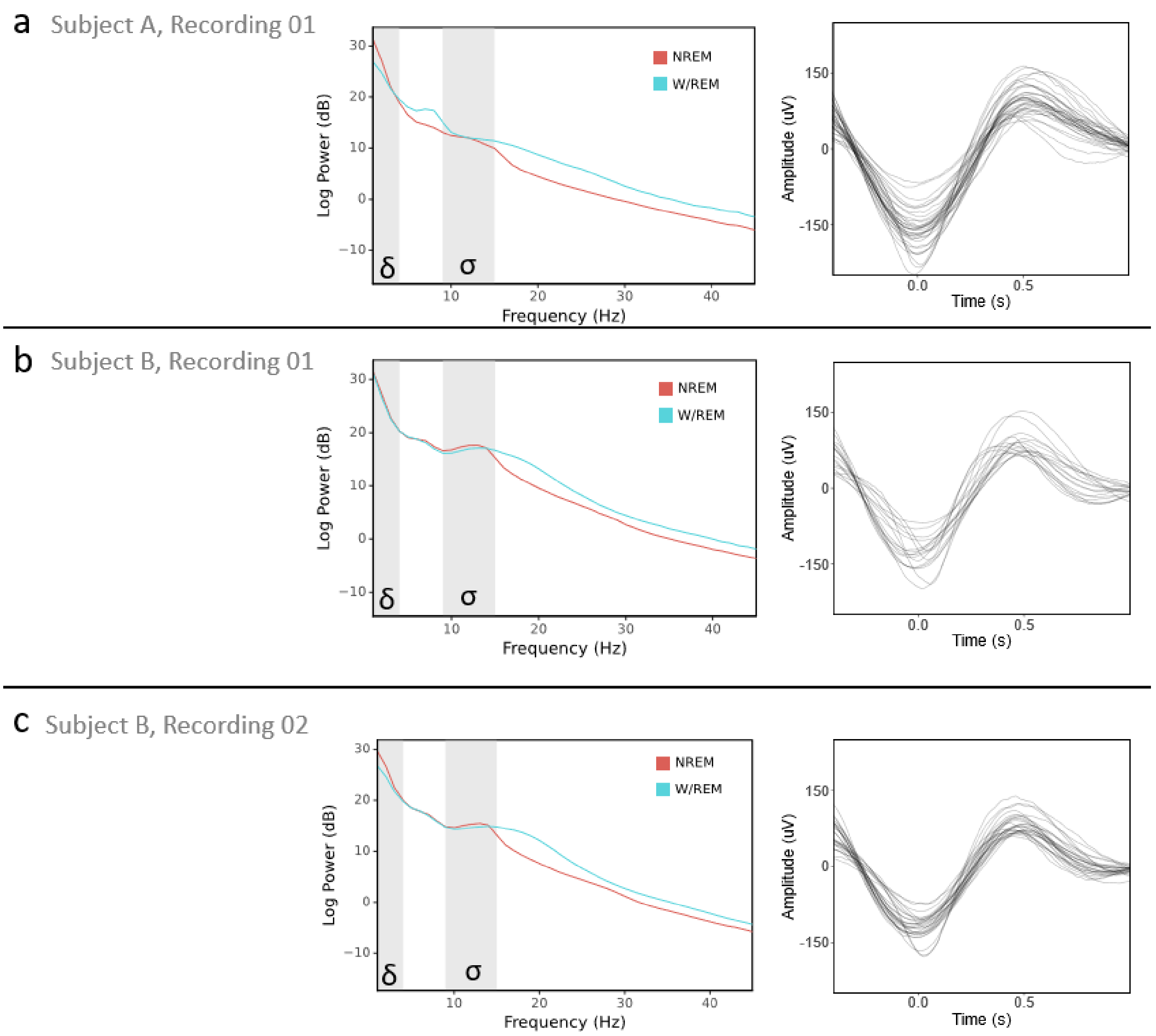


**Supplementary Figure 2. Power spectral densities and slow-wave waveforms.** (**a**) Subject A, Night 01, (**b**) Subject B, Night 01, and (**c**) Subject B, Night 02. Power spectral density plots of NREM (red) versus W/REM (blue) sleep across all macroelectrode channels (left panel). Delta and sigma frequency bands indicating SWA and sleep spindles, respectively (gray shading). Average waveforms of detected slow waves by macroelectrode channel (right).

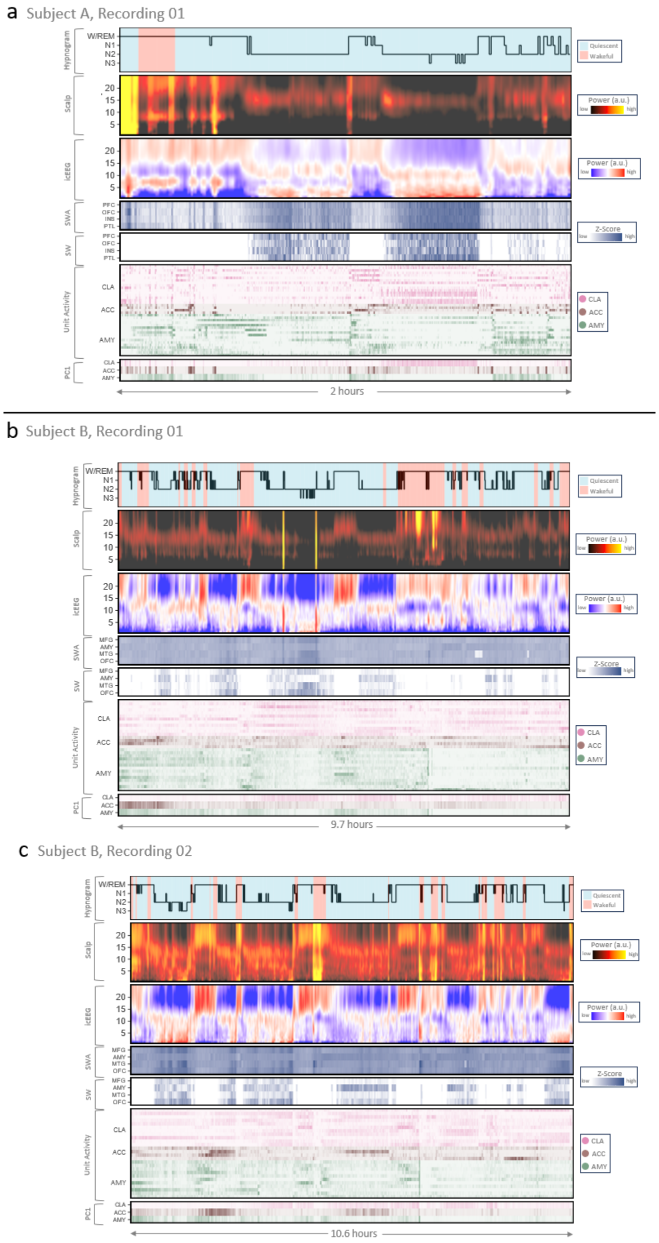


**Supplementary Figure 3. Sleep recording overviews.** (**a**) Subject A, Night 01, (**b**) Subject B, Night 01, and (**c**) Subject B, Night 02. Hypnogram shaded by behavioral state observed on avEEG: red indicates wakefulness and blue indicates behavioral quiescence (top row). Power spectrogram from the C4 scalp electrode (second row). Illustrative power spectrogram from the right middle frontal gyrus (third row). Binned z-score of slow-wave activity from four regions: middle frontal gyrus, amygdala, middle temporal gyrus, and orbitofrontal cortex (fourth row). Binned z-score of slow wave presence from the same regions (fifth row). Binned z-score of the firing rate for claustrum (pink), anterior cingulate cortex (brown), and amygdala (green) single units (sixth row). First principal component of the firing rate in the single units of the same regions (seventh row).

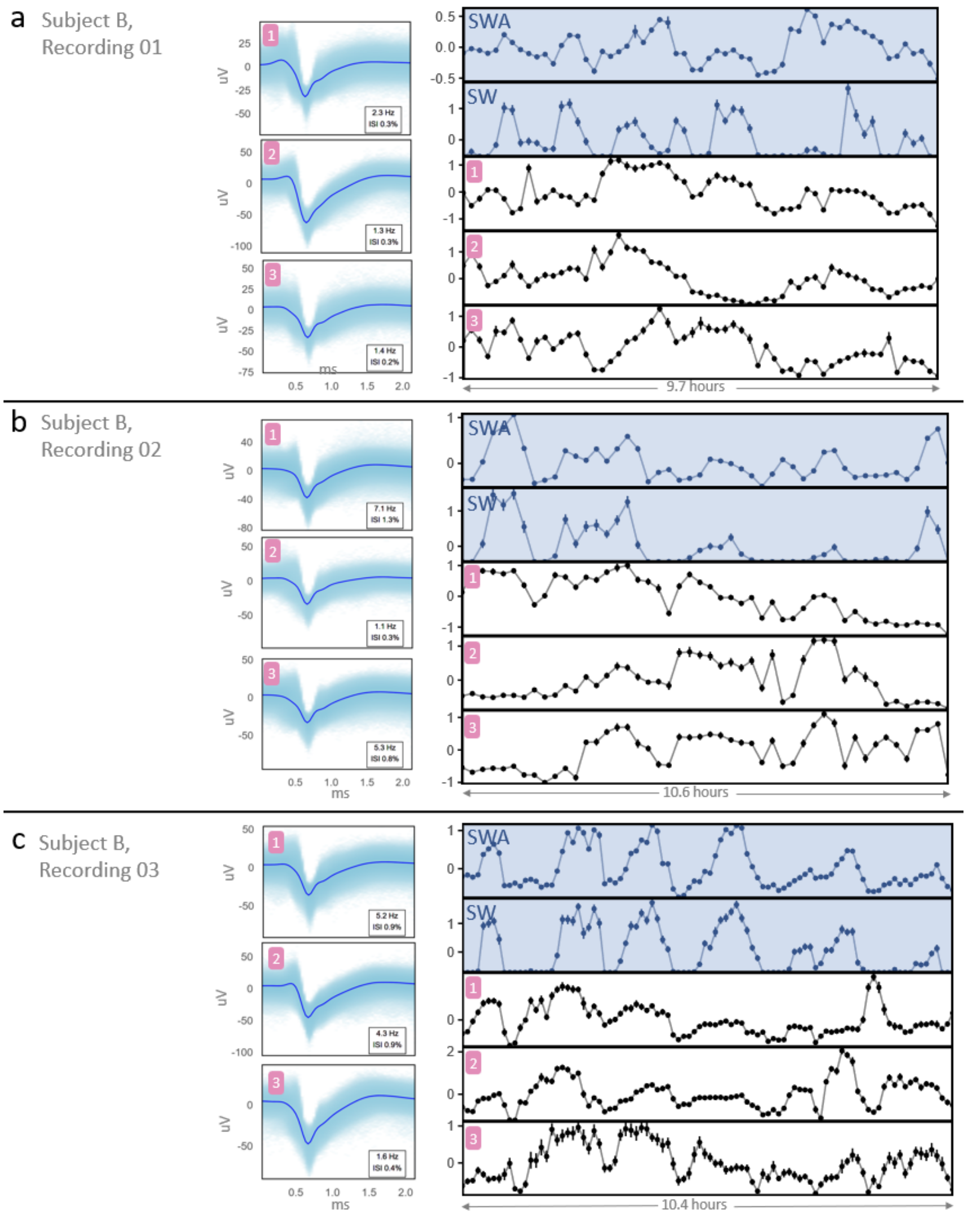


**Supplementary Figure 4. Example claustrum single units tracking SWA and SWs.** (**a**) Subject B, Night 01, (**b**) Subject B, Night 02, and (**c**) Subject B, Night 03. Average waveforms and inset unit statistics (firing rate and inter-spike interval violations) of three example claustrum single units for each sleep recording (left panel). Corresponding binned time series of z-scored firing rates (black lines) of the units aligned with the binned z-scored time series of representative slow-wave activity and slow wave presence (blue dots) in the right orbitofrontal cortex (right panel). Standard error is indicated by vertical bars.

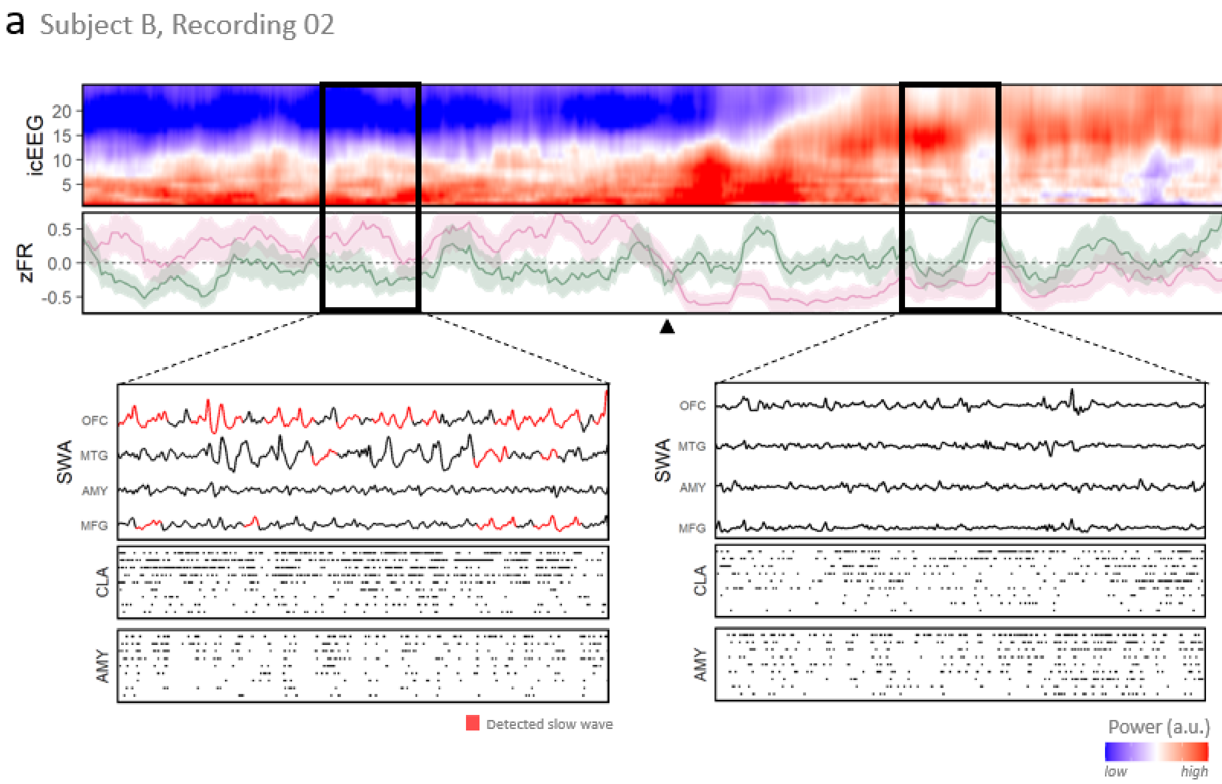


**Supplementary Figure 5. Transition out of NREM sleep in Subject B, Night 02.** Power spectrogram of left middle frontal macroelectrode centered on the time of sleep transition (first row). Z-score of population firing rates in the claustrum (pink) and amygdala (green) (second row). Magnified time windows before and after the sleep transition (bottom left and bottom right panels, respectively) containing time series representations of slow-wave activity for intracranial macroelectrode channels (red highlighting indicates a detected slow wave) in the orbitofrontal cortex, middle temporal gyrus, amygdala, and middle frontal gyrus (first rows of bottom panels) and single unit raster plots of spiking activity of the claustrum (second rows of bottom panels) and amygdala (third rows of bottom panels).

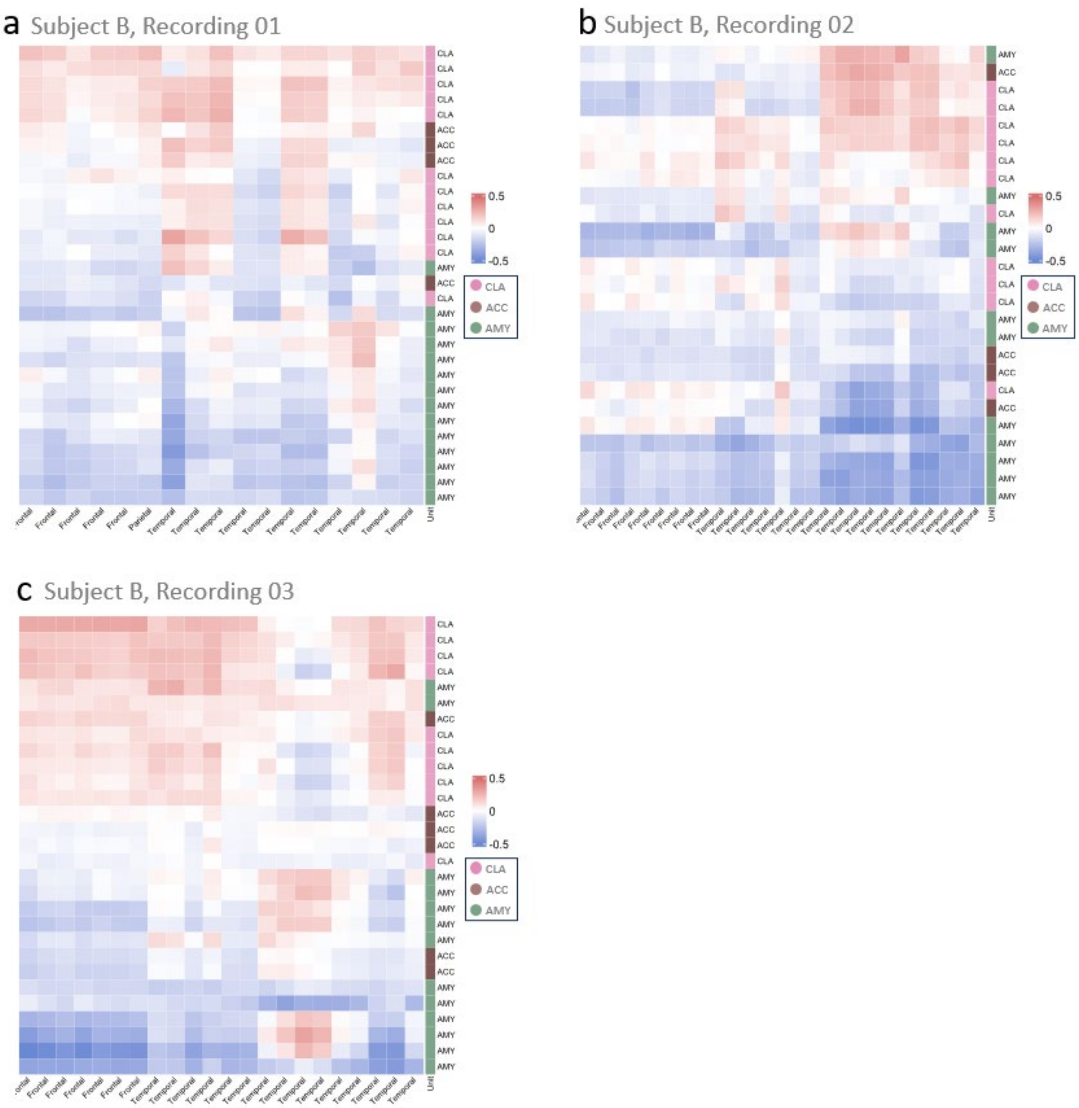


**Supplementary Figure 6. Heatmap of Spearman's ρ correlations of unit-channel pairs.** (**a**) Subject B, Night 01, (**b**) Subject B, Night 02, and (**c**) Subject B, Night 03. Single units are represented by rows and slow wave activity in macroelectrode channels are represented by columns. Colors indicate claustrum (pink), anterior cingulate (brown), and amygdala (green) single units. Each cell is colored according to the strength of the Spearman's ρ correlation.

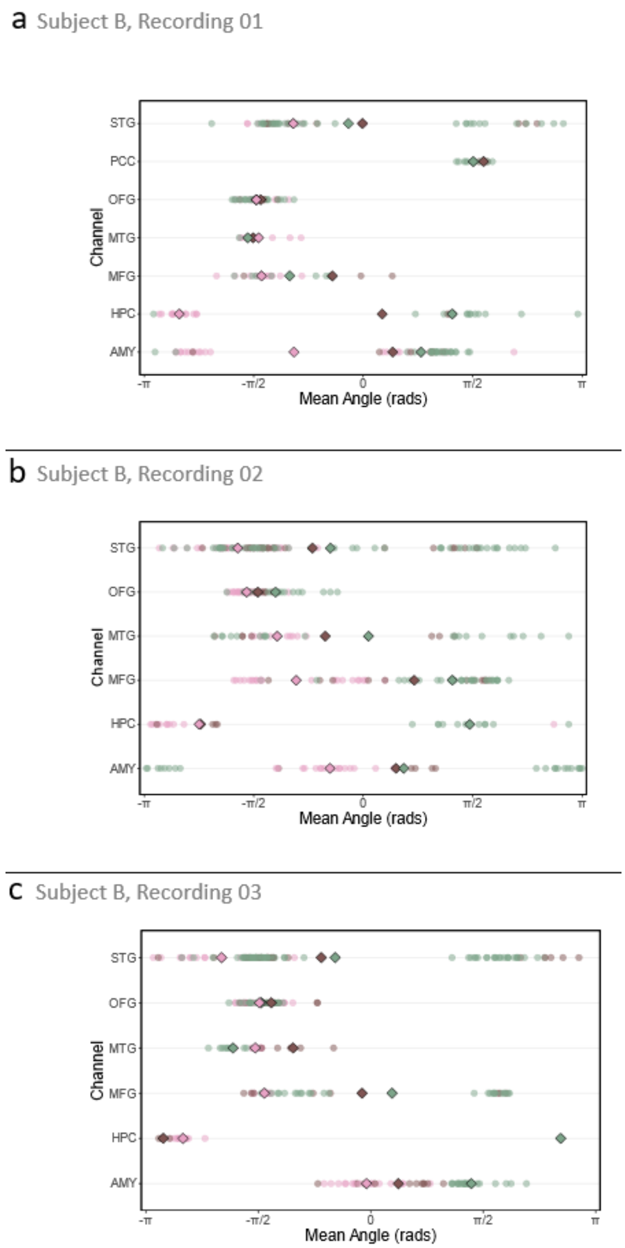


**Supplementary Figure 7. Spike-phase coupling.** Distribution of average phase angles for all unit-channel pairs with a phase distribution significantly different from uniform on Rayleigh's test. Results are organized by region of interest. **(a)** Subject B, Night 01, **(b)** Subject B, Night 02, and **(c)** Subject B, Night 03. Large diamonds indicate the average of all preferred phase angles for each unit region. Color indicates unit region.

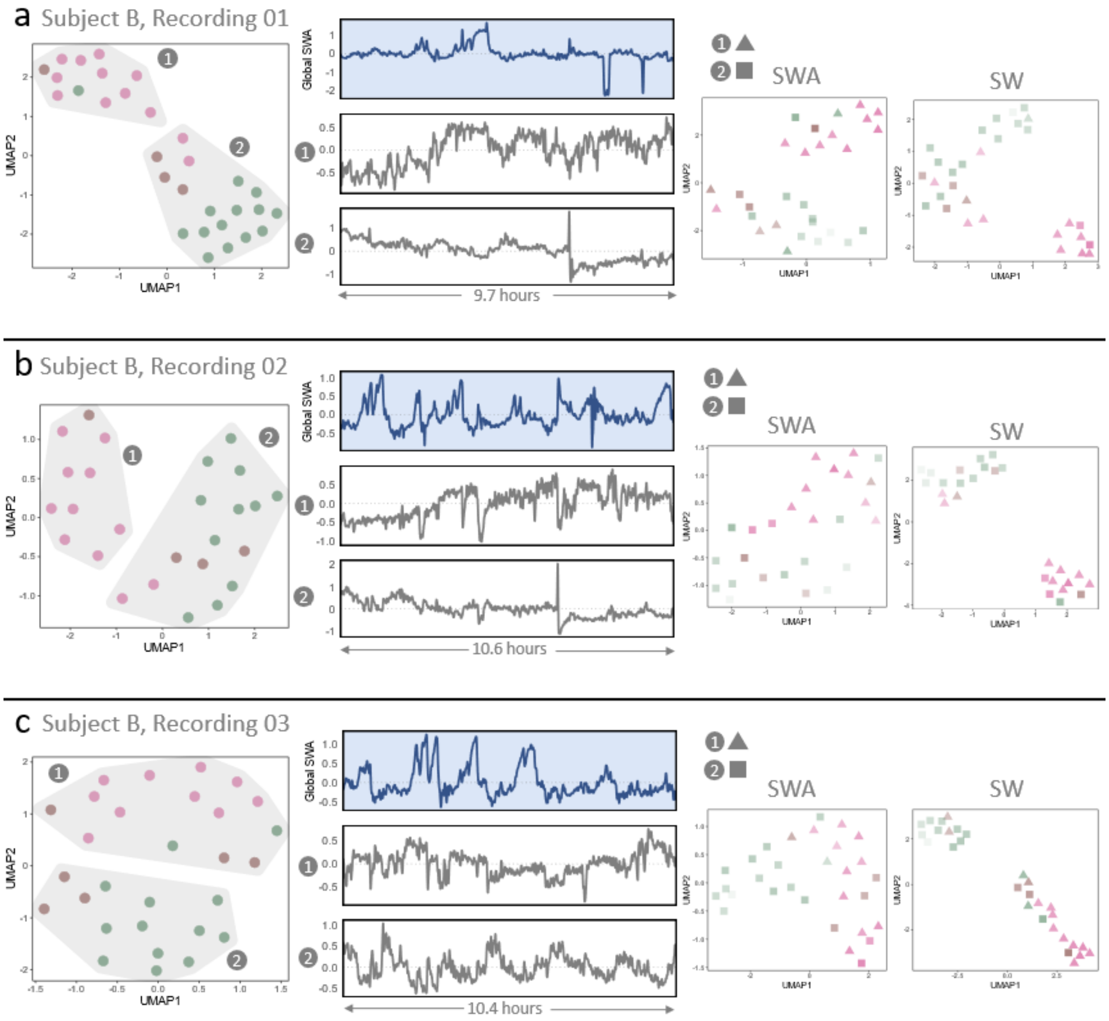


**Supplementary Figure 8. Claustrum spiking and correlation separates from controls after dimensionality reduction.** (**a**) Subject B, Night 01, (**b**) Subject B, Night 02, and (**c**) Subject B, Night 03. Scatterplot of two UMAP dimensions for single unit spiking activity demonstrates segregation of claustrum (pink), anterior cingulate (brown), and amygdala (green) units into two groups (labeled 1 and 2) indicated by gray shading (first panel). Time series of global slow-wave activity (top row, second panel) with aligned time series representing z-scored population firing rates for Group 1 (middle row, second panel) and Group 2 (bottom row, second panel). Scatterplot of two UMAP dimensions after dimensionality reduction of correlation with slow wave activity (third panel) and slow-wave presence (fourth panel) across channels. Group 1 units are indicated by triangles, and Group 2 units are indicated by squares.

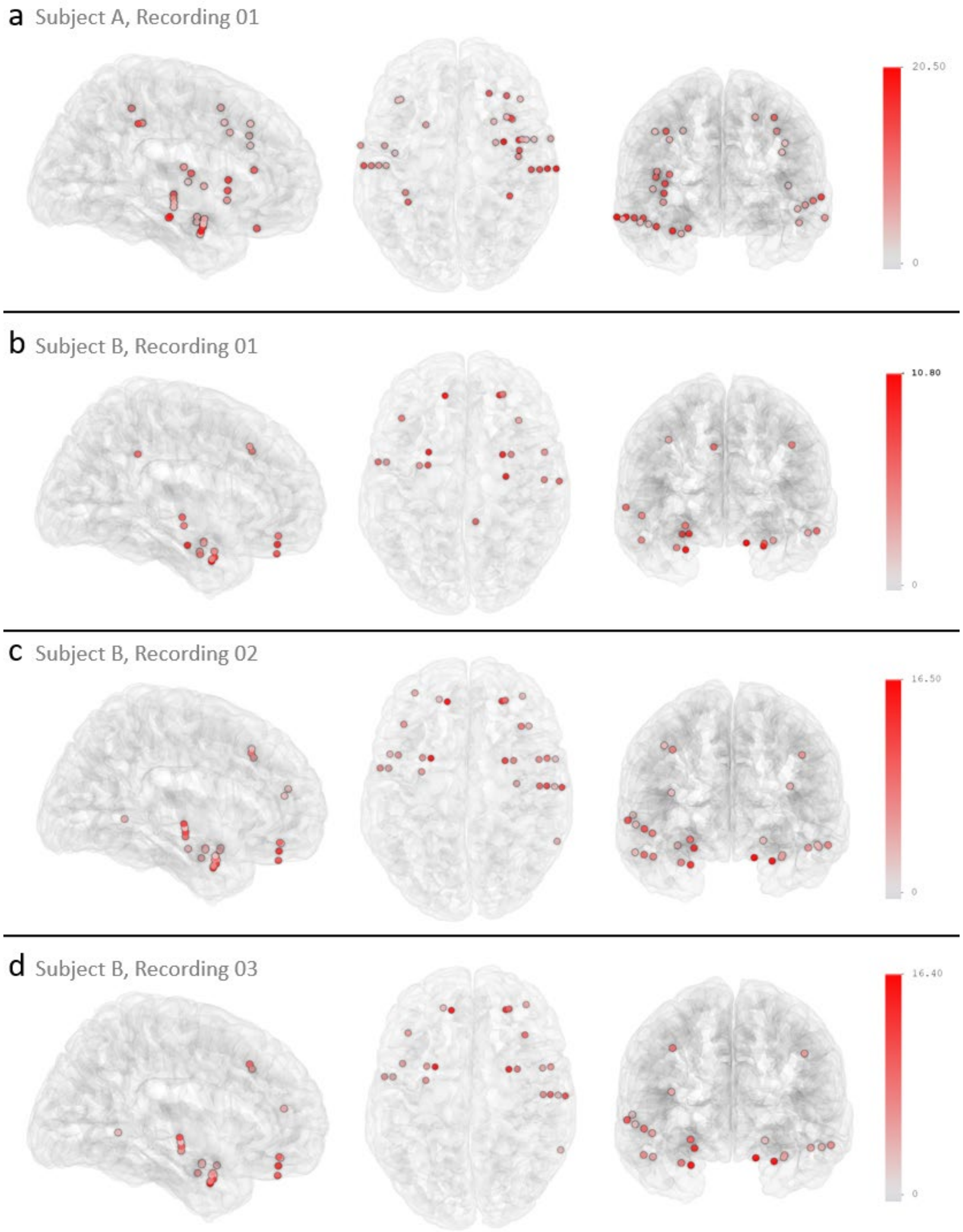


**Supplementary Figure 9. Detected slow waves per macroelectrode contact. (a)** Subject A, Night 01, **(b)** Subject B, Night 01, (**c**) Subject B, Night 02, **(d)** Subject B, Night 03. Right sagittal (left panels), top-down axial (middle panels), and anterior coronal (right panels) views of N27 brain template with superimposed dots indicating positions of macroelectrode contacts (after exclusion of under-detecting channels). Color indicates the average number of slow waves detected per minute (after slow-wave quality control).

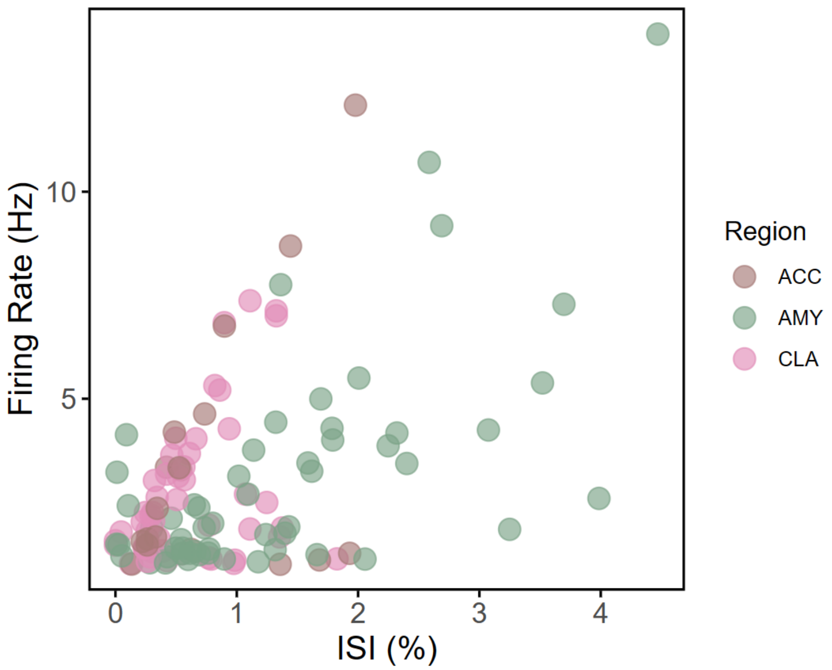


**Supplementary Figure 10. Spike sorting metrics for single units.** Scatterplot of the percent of inter-spike interval (ISI) violations < 3 ms (x-axis) and average firing rate (y-axis). Units are colored by region (brown = anterior cingulate cortex, green = amygdala, pink = claustrum). No CLA units had > 2% ISI violations, and only one AMY unit had > 4% ISI violations.
